# Machine learning–guided engineering of conditional split inteins for regulated protein splicing in mammalian cells

**DOI:** 10.1101/2025.10.21.683710

**Authors:** Junsheng Liang, Barbara Di Ventura

## Abstract

Inteins are proteins that excise themselves from precursor proteins and connect the flanking polypeptides with a peptide bond. Split inteins consist of two independently translated fragments that must associate to become splice-competent. They can be used for diverse post-translational protein modifications. Using the ML Int&in algorithm, we predicted unnatural split sites in two of the fastest and most efficient split inteins, gp41-1 and NrdJ-1, to generate functional variants with fragments of reduced mutual affinity. We harness this feature to create conditional versions of these inteins by controlling the physical proximity of the intein fragments with a light-inducible heterodimerization system. The resulting light-activatable gp41-1 and NrdJ-1 inteins enabled blue light–dependent control of Cre recombinase activity in mammalian cells, which we exploited to spatially control apoptosis via localized expression of truncated BID (tBID) and caspase-8. This work highlights the versatility of Int&in for designing conditional inteins for precise spatiotemporal protein regulation.

## Introduction

Inteins are short protein sequences embedded within host proteins that catalyze a self-excision process known as protein splicing, during which the intein removes itself and joins the surrounding flanking sequences (exteins) via a new peptide bond^1^. In nature, inteins are either encoded by a single gene giving rise to so-called contiguous inteins, which perform *cis*-splicing, or by two distinct genes, giving rise to so-called split inteins, which perform *trans*-splicing. In this case, the two fragments—termed N-intein and C-intein—must first associate to form a functional complex, after which the splicing reaction proceeds, ligating the N- and C-terminal exteins. Split inteins are especially valuable due to their broad utility in cell and synthetic biology^2-10^. Contiguous inteins have therefore been artificially split to generate new split inteins^11, 12^. Alternatively, they have been split at specific sites to obtain fragment pairs with low inherent affinity, which can be conditionally activated by bringing them into close proximity by either constitutive^13^ or inducible protein-protein interactions^11, 14, 15^. In all these cases, the decision where to split the intein was taken either heuristically or on the basis of simple structural assessments. We recently developed Int&in, a machine learning (ML) algorithm that predicts active split sites based on a combination of sequence and structural features, both local and global relative to the potential split position^16^. Interestingly, the model, based on logistic regression, demonstrates predictive power regarding splicing efficiency. Since the affinity between the intein fragments is a critical factor for the splicing reaction, we hypothesised that sites predicted to result in low-efficiency split inteins would most likely lead to two intein fragments with low binding affinity. These split inteins could then splice robustly if the two fragments are brought into physical proximity (Fig. 1a). The interaction between the intein fragments could be either constitutive ––e.g., using leucine zippers–– or inducible. This last case would allow the creation of conditional split inteins^1^.

**Fig. 1.**
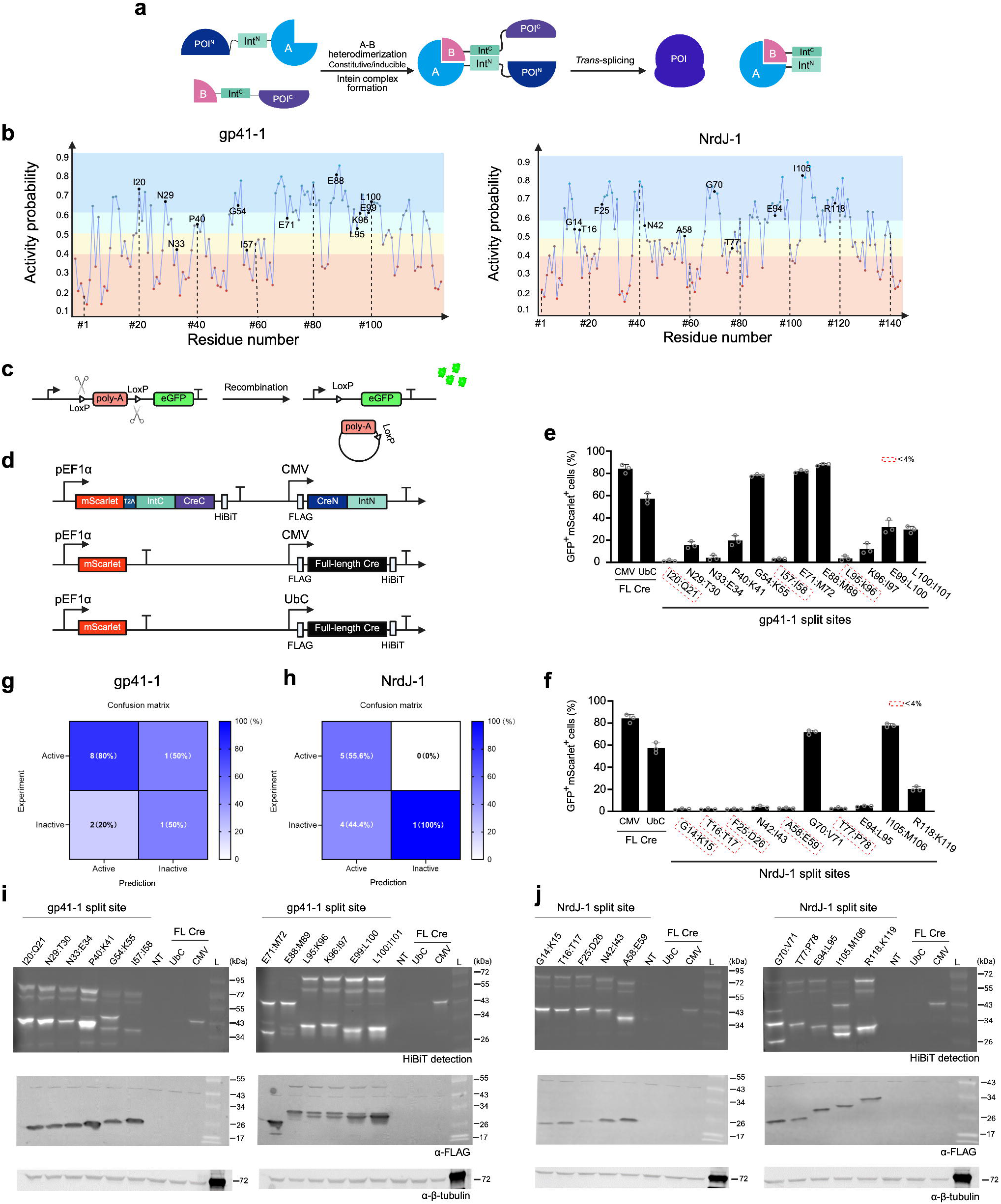
Using the ML algorithm Int&in to predict split sites in inteins that could be harnessed to engineer conditional inteins. **a** Schematic illustration of how a split intein could be regulated using a constitutively active or trigger-responsive heterodimerization system. Key to this approach is that the intein fragments (IntN and IntC) have low affinity for each other, so that their interaction is controlled by the heterodimerization system (A-B). POI, protein of interest. POI^N^, POI^C^, N- and C-terminal fragments into which the POI is split. The elements are not drawn to scale. **b** Output of the Int&in algorithm for the indicated inteins. The four categories are: active split sites (p ≥ 0.5), inactive split sites (p < 0.5), active with high probability (p ≥ 0.6), inactive with high probability (p < 0.4). The sites selected for experimental testing are indicated in black. Numbering starts at the fourth amino acid, as the first three residues correspond to extein sequences present in the crystal structure. Int&in is accessible at https://intein.biologie.uni-freiburg.de. **c** Schematic representation of the Cre-responsive reporter construct. **d** Schematic representation of the Cre constructs. **e, f** Bar graphs showing the percentage of cells transfected with the indicated constructs (mScarlet-positive) in which Cre successfully recombined the reporter DNA (GFP-positive) measured via flow cytometry. Red dotted squares indicate constructs which were labelled as inactive according to the selected threshold of 4% double-positive cells. Values represent mean ± SD of n = 3 biologically independent experiments. Individual data points are shown as open circles. FL Cre, full-length Cre expressed from either a strong (CMV) or weak (UbC) constitutive promoter. **g, h** Confusion matrix illustrating the performance of the Int&in prediction algorithm. Columns correspond to predicted classes, and rows to experimentally-determined classes. Percentages are normalized by column, indicating the proportion of correct predictions within each class. The intein was considered active if the percentage of double-positive cells was above 4%. **i, j** Representative Western blots of HEK 293 T cells transiently transfected with the indicated constructs performed to assess intein-mediated peptide bond formation. FL Cre, full-length Cre expressed from either a strong (CMV) or a weak (UbC) promoter. The N-terminal constructs as well as the splice product are detected with the anti-FLAG antibody. The C-terminal constructs as well as the splice product are detected via bioluminescence using the HiBiT tag. β-tubulin was used as loading control. NT, non-transfected cells. The experiment was repeated three times. Created with BioRender.

Here, we leverage the capability of Int&in to identify low-efficiency split sites to engineer blue light-inducible versions of two extremely fast and efficient split inteins, gp41-1 and NrdJ-1^17, 18^, to control the Cre recombinase. We optimize light-induced splicing by selecting the best heterodimerization system, mutating position -1 in the N-extein and by adding nuclear localization signals or nuclear export sequences to the constructs. We show that the optogenetic Cre we developed can be used to trigger apoptosis in selected cell sub-populations. These results show that Int&in is a valuable resource for engineering conditional split inteins, bypassing the trial-and-error phase typically required to identify active split sites, so that researchers can instead focus their efforts on the necessary optimization steps to achieve robust induction with fold-change and low leakiness for the specific protein of interest.

## Results

### Using the Int&in ML algorithm to predict split sites in gp41-1 and NrdJ-1 for conditional intein design

We started by selecting the split inteins to be made conditional. We chose the ultra-fast and efficient gp41-1 and the similarly fast and efficient NrdJ-1, both of which have been identified through bioinformatics analysis of metagenomic data collected from environmental marine microbial samples^18^ and later thoroughly characterized *in vitro*^17^ and applied in various cellular contexts including mammalian cells^2, 8, 9, 15, 19^. We fed the crystal structure of gp41-1^20^ (PDB ID: 6QAZ) and an AlphaFold2^21^-predicted structure of NrdJ-1 to the Int&in webserver^16^. We selected mostly sites with probabilities higher than 0.5 but below 0.7, corresponding to those predicted to be active with moderate splicing efficiency^16^. As mentioned above, we hypothesized that these sites would require a heterodimerization system to splice efficiently, making them ideal candidates for engineering conditional inteins. We also included a few sites with probabilities between 0.4 and 0.5 (Fig. 1b). Sites with probabilities below 0.4 are more likely truly inactive, as in this range the ratio of true negatives to false negatives is the highest^16^. Specifically, we selected 12 sites for gp41-1 (10 predicted to be active, and 2 to be inactive) and 10 sites for NrdJ-1 (8 predicted to be active, and 2 to be inactive). We avoided sites too close to the termini of the inteins, since previous work showed that splitting near the termini reduces splicing efficiency^22^. As a control, we included the natural split sites for both inteins (E88:M89 for gp41-1 and I105:M106 for NrdJ-1), which correspond to intein fragments with high affinity for each other, and are therefore expected to prevent external control over the splicing reaction.

### Engineering light-inducible Cre using the novel conditional split inteins

Next, we had to select the exteins. Instead of beginning with the reconstitution of model or “toy” fusion proteins—such as fluorescent proteins or other soluble, non-functional constructs—we chose to directly apply our engineering strategy to a functional protein. This decision was motivated by the consideration that each protein of interest typically requires its own optimization strategy. We selected the Cre recombinase as our target due to its broad utility and established role in synthetic biology^23^. To reconstitute Cre with split inteins, we first needed to identify a splice site within the protein that would generate two nonfunctional fragments, yet still support efficient protein *trans*-splicing by gp41-1 and NrdJ-1. Here, we use the term *splice site* to refer to a position intended for protein reconstitution through intein-mediated splicing and peptide bond formation, whereas *split site* denotes a position used to express a protein as two separate polypeptides that are not covalently rejoined. Both gp41-1 and NrdJ-1 require a serine at position +1—that is, the first residue of the C-extein, corresponding in this case to the C-terminal Cre fragment. We employed a previously identified splice site in Cre meeting these requirements^24^ (D109:S110), and additionally confirmed that each Cre fragment individually has no recombination activity (Supplementary Fig. 1). First, we had to test if gp41-1 and NrdJ-1 split at the selected unnatural sites retain activity on their own, when no heterodimerizer is included in the design. Sites for which splicing efficiently occurs without addition of the control element (in our case, the light-inducible heterodimerizer) cannot be employed to create conditional inteins. We set up an assay to assess Cre activity using a reporter construct in which a transcriptional terminator, placed upstream of the eGFP coding sequence, is flanked by the LoxP sites recognized by Cre (Fig. 1c). When transiently transfected in HEK cells alone, the reporter plasmid produces no fluorescence, confirming that there is no spontaneous recombination in the absence of Cre (Supplementary Fig. 2). Upon Cre-mediated recombination, the transcriptional terminator is excised and eGFP is expressed (Fig. 1c). We also cloned two positive controls: full-length Cre expressed under either the strong constitutive CMV promoter or the weaker constitutive UbC promoter^25^. The constructs included the fluorescent protein mScarlet under the strong constitutive EF1α promoter to report on successful transient transfection. We additionally included tags (FLAG, HiBiT) for Western blotting (Fig. 1d). We transiently transfected HEK 293T cells with the constructs and performed flow cytometry to quantify the percentage of cells positive for both red and green fluorescence (Fig. 1e,f), as well as the mean GFP fluorescence among the double-positive cells (Supplementary Fig. 3). A threshold of 4% double-positive cells was used to classify split sites as active. We found excellent agreement between the predictions and the experiments for gp41-1, with a precision of 80% (Fig. 1g). For NrdJ-1, Int&in performed less well, with a precision of 55.6% (Fig. 1h).

To confirm that *trans*-splicing occurred—and that fluorescence was not simply due to functional protein reconstitution without peptide bond formation—we performed Western blotting. A clear splice product was detected for the inteins split at sites corresponding to high fluorescence in the flow cytometry experiment (Fig. 1i,j). Interestingly, full-length Cre driven by the weak UbC promoter was also practically undetectable on the Western blot despite being only circa 30% less active than CMV-driven Cre (Fig. 1i,j). To further prove that *trans*-splicing occurs, we mutated the inteins at the critical asparagine (last residue of the intein) and cysteine (first residue of the intein) and performed flow cytometry. The percentage of double-positive cells was drastically reduced to less than 4% for all the sites but E99:L100 and L100:I101 for gp41-1 and I105:M106 and R118:K199 for NrdJ-1 (Supplementary Fig. 4). We concluded that the GFP fluorescence is the result of intein-mediated protein splicing.

Next, we turned to the selection of an inducible heterodimerizer. We opted for a light-inducible one because light is an excellent external trigger, which can be applied with high spatiotemporal precision in a non-invasive way. We chose iLID^26, 27^, a widely used system known for its robustness. The two interacting partners (iLID and SspB_nano) were cloned so as not be included in the splice product (Fig. 2a, upper panel). We tested all the gp41-1 split sites except G54:K55 and E71:M72, which had resulted in almost as high Cre activity as the natural split site E88:M89 in the absence of any heterodimerizer (Fig. 1d). These sites are likely to be non-controllable. As a proof of this, we included the natural E88:M89 split site in the design with iLID. We also included a control in which only the light-inducible heterodimerizer was present, but no intein (Fig. 2a, lower panel). This control would allow us to distinguish the contribution of *trans*-splicing from that of protein-protein interactions in the Cre activity assay. Cloning of the construct bearing the I57:I58 split site was unsuccessful. As this site was predicted and experimentally confirmed to be inactive in the screen without the heterodimerizer (Fig. 1i), no further attempts were made to recover it.

**Fig. 2.**
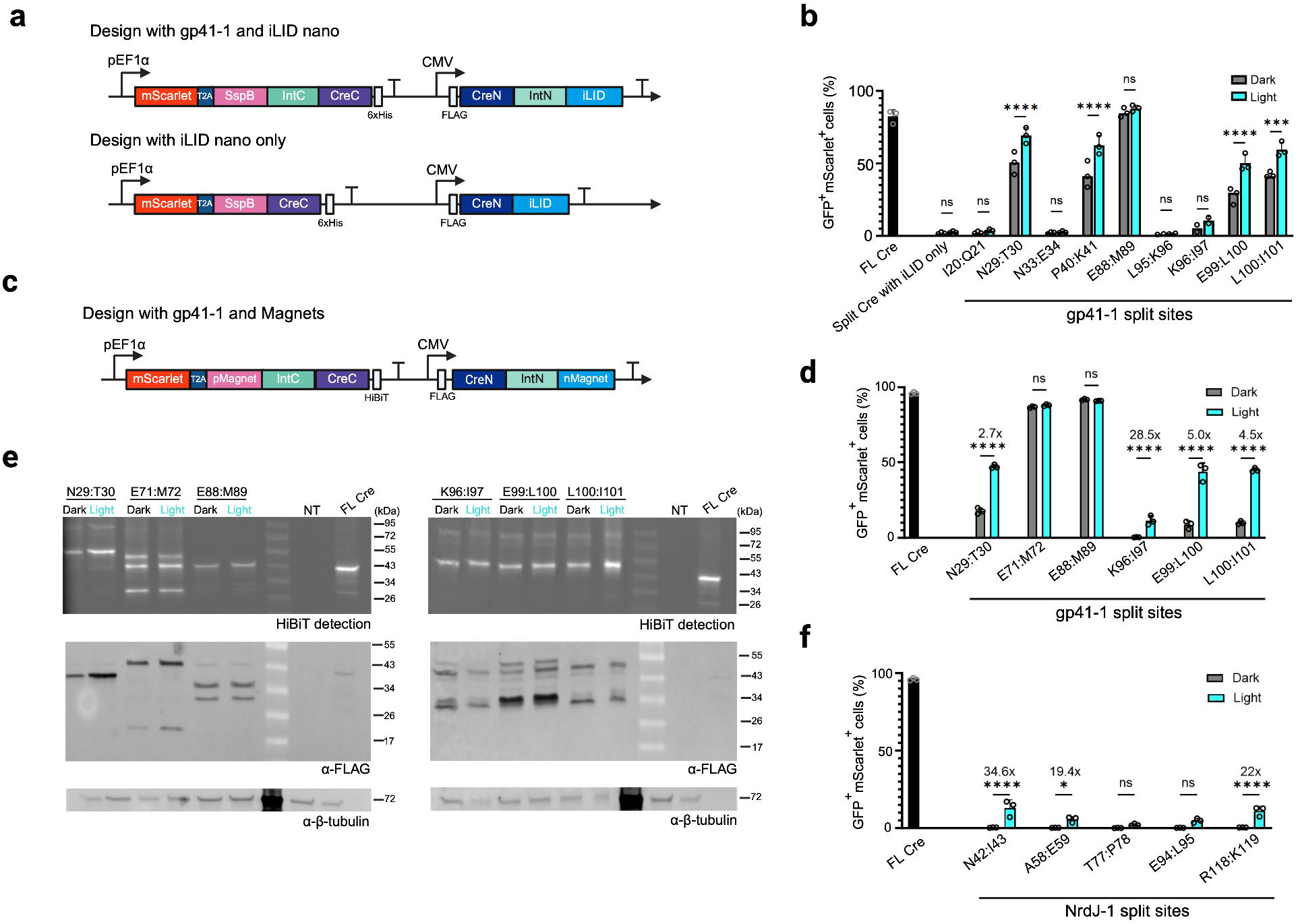
Engineering conditional gp41-1 and NrdJ-1 with the blue light-responsive Magnets heterodimerization system. **a**,**c** Schematic representation of the constructs. T2A, ribosome skipping sequence of the Thosea asigna virus. **b**,**d**,**f** Bar graphs showing the percentage of HEK 293T cells transfected with the indicated constructs (mScarlet-positive) in which Cre successfully recombined the reporter DNA (GFP-positive) measured via flow cytometry. Values represent mean ± SD of n = 3 biologically independent experiments. Individual data points are shown as open circles. FL Cre, full-length Cre expressed from the strong, constitutive CMV promoter. Cells were either kept in the dark the whole time or first illuminated in pulsed cycles of 20 s blue light (13 mW/cm^2^) followed by 60 s dark, repeated for 6 hours, then kept in the dark for 18 hours before the measurement. Cells expressing FL Cre were kept in the dark. Statistical significance was determined by unpaired, two-tailed Student’s *t*-tests, with multiple-comparison correction according to the Benjamini–Krieger–Yekutieli procedure. ****, p-value ≤ 0.0001; ***, p-value ≤ 0.001; *, p-value ≤ 0.05; ns, p-value > 0.05. **b** Light-dark fold change is indicated for some of the constructs. (**d**,**f**) Light-dark fold change is indicated for the constructs with statistically significant difference between light and dark values. Cells transiently transfected with full-length Cre were kept in the dark. **e**, Western blot of HEK 293T cells transiently transfected with the indicated constructs performed to assess intein-mediated peptide bond formation. Cells were treated as in (**b**,**d**,**f**), except cells were lysed 2 hours post-illumination. FL Cre, full-length Cre, expressed from the strong, constitutive CMV promoter. The N-terminal constructs as well as the splice product are detected with the anti-FLAG antibody. The C-terminal constructs as well as the splice product are detected via bioluminescence using the HiBiT tag. β-tubulin was used as loading control. NT, non-transfected cells. Created with BioRender.

It is known that the amount of transfected DNA influences the behaviour of synthetic constructs, particularly optogenetic ones, with higher expression levels typically leading to higher leakiness. To minimize splicing activity in the dark, cells were transfected with low amounts of construct DNA (50 ng per well, with 10^5^ cells seeded per well). The total amount of transfected DNA was kept constant by supplementing with stuffer plasmid. We either kept the cells in the dark for 24 hours or first illuminated them with pulsatile blue light for 6 hours and then incubated them back in the dark for additional 18 hours to let the reporter accumulate. We then performed flow cytometry to quantify the number of cells positive for both GFP (Cre reporter) and mScarlet (transfection control). We found some light inducibility for 5/6 sites that were labelled as active according to the data without the heterodimerizer, albeit not statistically significant (Fig. 2b). As expected, the natural split site did not allow for light control (Fig. 2b). For gp41-1 split at K96:I97, there was a trend, but no statistically significant difference between light and dark samples. For the I20:Q21 split site, there was no Cre activity, consistent with its lack of activity in the absence of the dimerizer (Fig. 1e). Interestingly, iLID on its own had no effect (Fig. 2b).

Despite having obtained light-mediated Cre activity for most split sites, we were not satisfied, as the background—that is, activity in the dark—was quite high in all cases but one (K96:I97). In an attempt to find a more optimal design, we tested the construct based on the N29:T30 split site–showing the highest light-induced activity–with alternative iLID variants (sLID and SspB_micro instead of SspB_nano)^27^, as well as with the addition of a nuclear localization signal (NLS) and/or a nuclear export sequence (NES) in various combinations (Supplementary Fig. 5, upper panel). None of these modifications significantly improved performance. One resulted in substantially lower leakiness, but this was accompanied by a marked reduced in light-induced activity (Supplementary Fig. 5, lower panel). Thus, we turned to a different heterodimerization system, namely Magnets^28^ (Fig. 2c). This system is also widely employed and known to be reliable. We cloned the same constructs as for iLID, leaving out the one bearing the N33:E34 split site which had very low activity on its own and almost no activity with iLID. We additionally included as a control the two unnatural split sites that resulted in very high Cre activity in the assay without heterodimerizer (G54:K55 and E71:M72; Fig. 1e). We also again included the control with only Magnets and no intein (Supplementary Fig. 6, upper and middle panels). We transiently transfected HEK 293T cells and performed flow cytometry. This time, we found much less Cre activity in the dark for all sites except for those we expected to be not controllable (Supplementary Fig. 6, lower panel, and Fig. 2d). Sites I20:Q21 and L95:K96 still showed no Cre activity, suggesting they are truly inactive. The construct without intein but only Magnets also showed no Cre activity (Supplementary Fig. 6, lower panel). The GFP mean intensity was also higher in the light than in the dark for all light-responsive constructs but one (Supplementary Figure 7). To confirm that Cre activity was a consequence of *trans*-splicing, we performed Western blotting. On the blot, there was a clear band running at the size of full-length Cre for the controls (natural split site E88:M89, and split site E71:M72); for the construct bearing the N29:T30 split site, a faint band could be detected in the illuminated sample, confirming the light-dependence of the reaction (Fig. 2e). For the other three constructs that gave a robust light-dark difference in Cre activity, we could not detect a band (Fig. 2e). However, this is also true for the full-length Cre whose expression is driven by the weaker UbC promoter (Fig. 1i,j). To further confirm that gp41-1’s splicing activity is necessary to regain Cre activity, we mutated the critical intein residues (conserved N-terminal cysteine and C-terminal asparagine) within the Magnets’ constructs. With the mutated gp41-1 we did not detect any Cre activity in the flow cytometer (Supplementary Fig. 8).

Given that Magnets outperformed iLID in the gp41-1 constructs, we went on to implement this heterodimerizer with NrdJ-1. As unnatural split sites, we selected all those that showed very little activity in the absence of the heterodimerizer (Fig. 1f). We transiently transfected HEK 293T cells with the constructs, and either kept the cells in the dark for 24 hours or illuminated them with pulsatile blue light for 6 hours followed by incubation back in the dark for an additional 18 hours to allow the reporter to accumulate. We then performed flow cytometry to quantify the number of cells positive for both GFP (Cre reporter) and mScarlet (transfection control). For three out of the five tested constructs, there was light-mediated Cre activity, albeit with a low percentage of positive cells (Fig. 2f). As for GFP intensity in the double-positive cells, we found a statistically significant difference between light and dark conditions for the constructs bearing the N42:I43, A58:E59 and R118:K119 split sites (Supplementary Fig. 9).

### Further optimization of the light-inducible Cre constructs

At first, we sought a scarless Cre reconstitution, splitting the protein at an appropriate serine residue that would support *trans*-splicing without having to mutate or insert amino acids into the enzyme. However, inteins are sensitive to the so-called local exteins, that is, the amino acids immediately flanking them, and each intein will optimally splice in the presence of specific local exteins, which might not necessarily work well with a different intein^29^. Searching for information on local extein preference for NrdJ-1, we found a recent study showing that an aspartate at position -1 is suboptimal^30^. To assess whether the local exteins limited the amount of splicing—and, consequently, the amount of Cre activity—obtained with our NrdJ-1 constructs, we constructed a series of mutants, either by mutating only position -1 according to the results of Dheer and Muir^30^ (Fig. 3a) or by adding the optimal six amino acids at positions -1, -2, -3 and +1, +2, and +3 (Fig. 3b). Since the splice site is located in a flexible linker of Cre, we hypothesized that the addition of these extra amino acids would have negligible impact on enzyme activity. We confirmed this hypothesis by transiently transfecting all the modified Cre constructs into HEK 293 cells and performing flow cytometry (Supplementary Fig. 10). We additionally included a design whereby, upon *trans*-splicing, the protein is fused to an N-terminal and a C-terminal NLS (Fig. 3a,b). We implemented the changes into two light-inducible NdrJ-1 constructs (N42:I43 and R118:K119; Fig. 2f). Moreover, we decided to transfect 150 ng instead of 50 ng of construct DNA (per well, with 10^5^ cells seeded per well) hoping this would improve the performance of the NrdJ-1 constructs. We found that even mutation of the -1 position alone (from D to C or W, respectively) had a large influence on the performance of the constructs: the percentage of double-positive cells after blue light exposure went from 14% to 43% and 25%, respectively, for those bearing NrdJ-1 split at position N42:I43 (Fig. 3c). The fold-change went from 23.6 for the original construct to 65 for the construct with Cre^D109W^. An improvement was seen also for the constructs bearing NrdJ-1 split at position R118:K119 (Fig. 3d). In particular, mutation of D to C led to a 3.5-fold increase in the percentage of double-positive cells in the light, albeit the leakiness also increased. For the construct based on NrdJ-1 split at position N42:I43, the addition of the six local exteins yielded double the activity compared to the original construct (Fig. 3c), while for the one based on NrdJ-1 split at position R118:K119, the improvement was modest (Fig. 3d). Since Cre is active in the nucleus, a more pronounced nuclear accumulation is expected to be beneficial. Indeed, the addition of the NLSs led to an improvement for many of the constructs (Fig. 3c,d). For two of the constructs with high activity in the light, we confirmed *trans*-splicing via Western blotting (Fig. 3e).

**Fig. 3.**
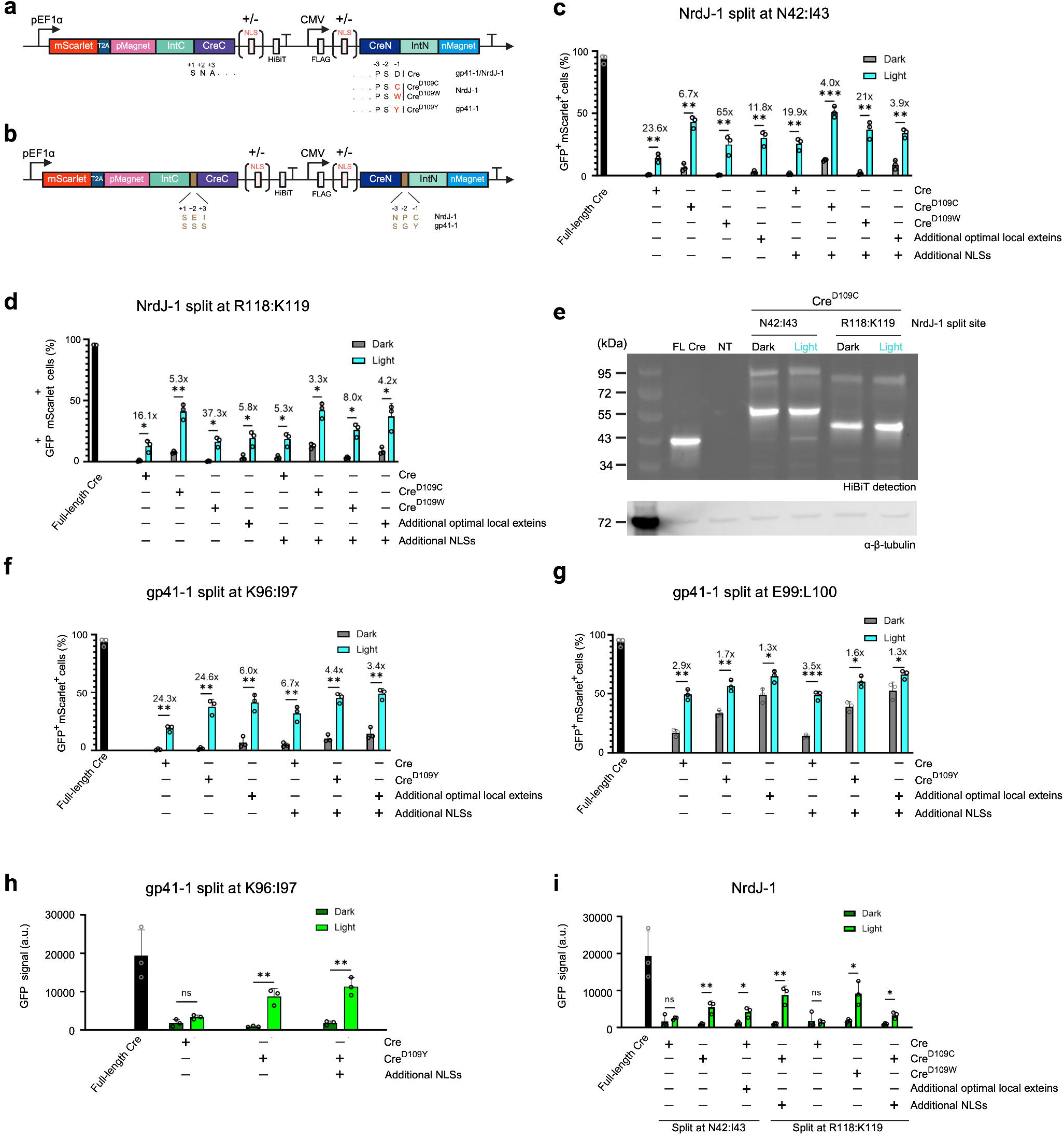
Construct optimization yields several high-performing light-inducible Cre variants. **a**,**b** Schematic representation of the constructs. The same constructs were made with (+) and without (-) additional nuclear localization signals (NLSs) on the N- and C-terminal constructs. T2A, ribosome skipping sequence of the Thosea asigna virus. **a**, Constructs with mutations in the last amino acid of the N-terminal Cre fragment, representing position -1 in respect to the intein. **b**, Constructs with extra amino acids in Cre. These are the optimal local exteins for the intein. **c**,**d**,**f**,**g**,**h**,**i** Bar graphs showing the percentage (**c**,**d**,**f**,**g**) or the mean GFP fluorescence (**h**,**i**) of cells transfected with the indicated constructs (mScarlet-positive) in which Cre successfully recombined the reporter DNA (GFP-positive) measured via flow cytometry. Values represent mean ± SD of n = 3 biologically independent experiments. Individual data points are shown as open circles. Full-length Cre was expressed from the strong constitutive CMV promoter. Cells were either kept in the dark the whole time or first illuminated in pulsed cycles of 20 s blue light (13 mW/cm^2^) followed by 60 s dark, repeated for 6 hours, then kept in the dark for 18 hours before the measurement. Cells expressing FL Cre were kept in the dark. Statistical significance was determined by unpaired, two-tailed Student’s *t*-tests, with multiple-comparison correction according to the Benjamini–Krieger–Yekutieli procedure. ****, p-value ≤0.0001; ***, p-value ≤ 0.001; **, p-value ≤0.01; *, p-value ≤ 0.05; ns, p-value > 0.05. The light-dark fold change is indicated for the constructs with statistically significant difference between light and dark values. **e** Western blot of HEK 293T cells transiently transfected with the indicated constructs performed to assess intein-mediated peptide bond formation. Cells were treated as in (**c**,**d**,**f**,**g**,**h**,**i**), except cells were lysed 2 hours post-illumination. FL Cre, full-length Cre, expressed from the strong, constitutive CMV promoter. Detection was performed via bioluminescence using the HiBiT tag. β-tubulin was used as loading control. NT, non-transfected cells. Created with BioRender.

Next, we assessed whether the same strategy could also improve the gp41-1-based constructs. Here we tested only a mutation at position -1 from D to Y, the addition of the optimal local exteins for gp41-1, and the designs with the two NLSs (Fig. 3a,b). For the construct bearing gp41-1 split at K96:I97, the simple mutation of position -1 within Cre from D to Y led to a 2-fold improvement in the amount of light-induced recombination (Fig. 3f). The addition of the six local exteins also had a substantial impact, although, in this case, the leakiness increased as well (Fig. 3f). This was true for all other tested constructs. Interestingly, for the construct bearing gp41-1 split at E99:L100, all modifications worsened the performance—as the leakiness dramatically increased—, with the exception of the construct bearing the NLSs, which was slightly improved in terms of tightness compared with the original one (Fig. 3g). We found a clear improvement also for the mean GFP signal in the double-positive cell population (Fig. 3h). To demonstrate the versatility of the system, we also tested some of the best performing constructs in U2OS cells. Although efficiencies were lower, we still observed light-inducibility in all cases (Supplementary Fig. 11).

### Steering Cre activity by modulating light exposure

The advantage of light is that it can be easily confined in space. To showcase the possibility of activating Cre in selected cell sub-populations, we shined blue light onto the cells through a photomask (Fig. 4a). The pattern is clearly visible (Fig. 4b), albeit not as striking as for those obtained with bacteria^31^ because mammalian cells cannot be plated at very high densities and also move during the experiment.

**Fig. 4.**
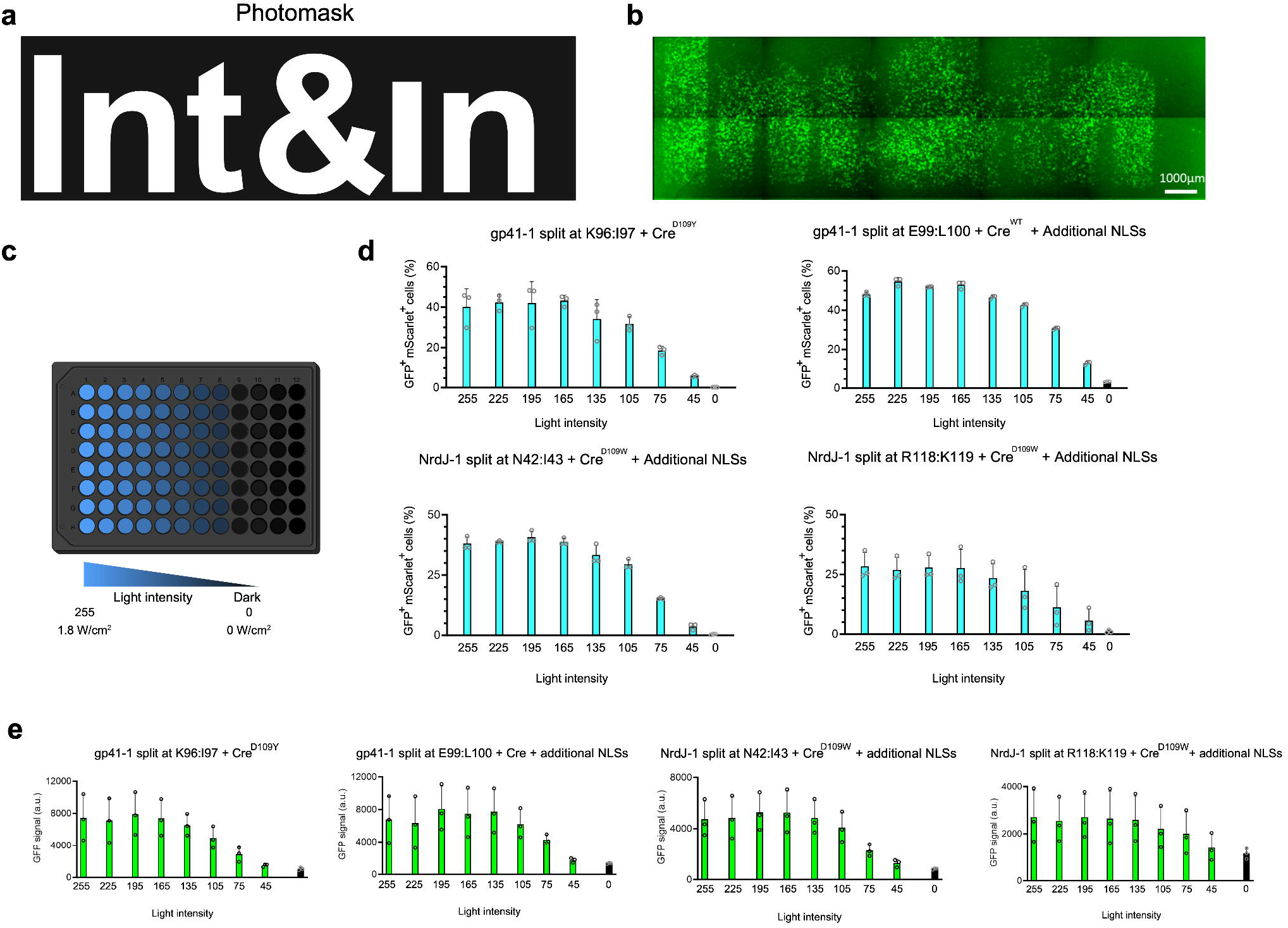
The splicing activity of the engineered inteins can be controlled in space and tuned via light intensity. **a** Representation of the photomask used to illuminate the cells. **b**, Fluorescence microscopy images of HEK 293T cells transiently transfected with the construct based on gp41-1 split at K96:I97 bearing Cre^D109Y^ illuminated for 2 hours with blue light (455 nm, 2.3 mW/cm^2^), then kept in the dark for 22 hours. **c** Schematic representation of the setup used to assess the effect of light intensity. A black 96-well plate was placed on the Diya illumination device 24 hours post-transfection. Each well is illuminated with an individual optical channel with blue light for 12 hours, followed by 12 hours in the dark before measurement. A device value of “255” corresponds to light intensity of 1.8 mW/cm^2^. **d**,**e** Bar graphs showing the percentage (**d**) or the mean GFP fluorescence (**e**) of HEK 293T cells transfected with the indicated constructs (mScarlet-positive) in which Cre successfully recombined the reporter DNA (GFP-positive) measured via flow cytometry. Values represent mean ± SD of n = 3 biologically independent experiments. Individual data points are shown as open circles.

Light intensity modulation represents another parameter to control the extent of Cre activity. To demonstrate this, we exposed HEK 293T cells transiently transfected with four selected constructs to a gradient of light intensities using the Diya illumination device^32^ (Fig. 4c). Cre activity was then quantified by flow cytometry. All constructs showed a clear dose-response relationship (Fig. 4 d,e). Notably, variability was higher in this experiment compared with others, likely due to greater differences in transfection efficiencies in the 96-well plate format.

### Triggering apoptosis with light by Cre-mediated expression of caspase-8 and tBID

So far, we used Cre to control the expression of the fluorescent protein eGFP. To showcase the utility of the developed light-inducible Cre, we decided to swap eGFP with a functional protein. Apoptosis is an essential cellular process particularly important during development^33^. Tissue engineering benefits from the possibility to trigger apoptosis in a spatially and temporally defined manner to create desired architectures or cavities^34^. For this reason, we decided to control apoptosis with our conditional intein system. A key player in both the intrinsic and extrinsic apoptotic pathways is BID (BH3 interacting domain death agonist)^35^. In the extrinsic pathway, BID is truncated into its active form, tBID, by caspase-8^36, 37^. In the intrinsic pathway, cathepsins–lysosomal proteases– have been shown to cleave and activate BID upon lysosomal rupture and escape into the cytosol^38^. Granzymes released by cytotoxic T lymphocytes and natural killer cells can likewise cleave BID to generate tBID^39, 40^. tBID then permeabilizes the mitochondrial outer membrane, either directly^41-43^ or via BAX/BAK^44, 45^, leading to the release of cytochrome c and eventually activation of effector caspases, such as caspase 3, which execute cell death^35^ (Fig. 5a). Given its role in both the intrinsic and extrinsic apoptotic pathways, we opted to control tBID, which does not need upstream processing by other proteins to function. We also selected caspase-8 as an additional candidate to control apoptosis. In this case, caspase-8 would act by cleaving endogenous BID into tBID. We replaced the *egfp* gene with the genes encoding these proteins in the Cre-responsive expression cassette (Fig. 5b). We transiently transfected HEK 293T cells with the constructs and either kept the cells in the dark for 24 hours or exposed them to blue light in a pulsed regime for 6 hours, followed by an additional 18 hours incubation in the dark. We quantified apoptosis by flow cytometry using Annexin V, a phospholipid-binding protein that recognizes phosphatidylserine exposed on the outer leaflet of the plasma membrane during early apoptosis, together with Helix NP, a nucleic acid stain impermeant to live cells. Annexin V-positive, Helix NP-negative cells were classified as early apoptotic; Annexin V-negative, Helix NP-positive cells as primarily necrotic; and Annexin V/Helix NP double-positive cells as late apoptotic or secondarily necrotic. We observed light-dependent apoptosis in both systems, with the tBID-based approach showing particularly low background activity (Fig. 5c).

**Fig. 5.**
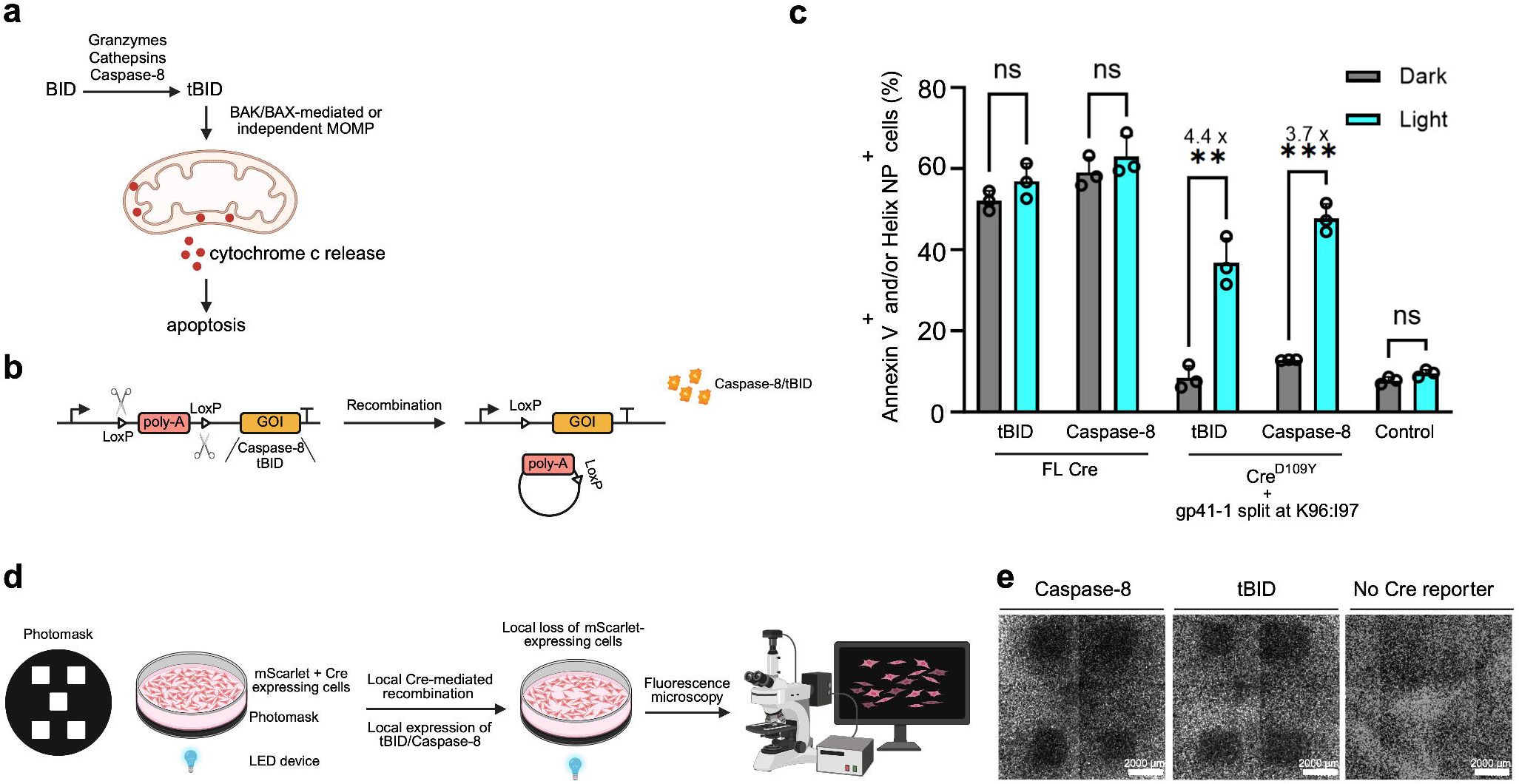
Controlling apoptosis with blue light. **a** Schematic drawing showing some of the key molecular species involved in the initiation of apoptosis in mammalian cells. **b** Schematic representation of the apoptosis-triggering constructs. **c** Bar graph showing the percentage of cells transfected with the indicated constructs that underwent apoptosis or necrosis. Quantification was performed via flow cytometry. Values represent mean ± SD of n = 3 biologically independent experiments. Individual data points are shown as open circles. Statistical significance was determined by unpaired, two-tailed Student’s *t*-tests, with multiple-comparison correction according to the Benjamini–Krieger–Yekutieli procedure. ****, p-value ≤0.0001; ***, p-value ≤ 0.001; **, p-value ≤0.01; *, p-value ≤ 0.05; ns, p-value > 0.05. The light-dark fold change is indicated for the constructs with statistically significant difference between light and dark values. FL Cre, full-length Cre expressed from the strong, constitutive CMV promoter. Cells were either kept in the dark the whole time or illuminated for 6 hours with blue light (2.3 mW/cm^2^) in short pulses (20 s light / 60 s dark) followed by additional incubation in the dark for 18 hours. Control, cells transfected only with stuffer plasmid DNA. **d** Schematic representation of the workflow to trigger apoptosis only in selected areas of the culture dish. **e** Representative fluorescence microscopy images acquired using the Texas Red channel on a Lionheart FX automated microscope at 1.25x magnification. HEK 293T cells were illuminated for 6 hours with blue light (2.3 mW/cm^2^) in short pulses (20 s light / 60 s dark) followed by additional incubation in the dark for 24 hours. Scale bar, 2000 μm. Created with BioRender.

In this experiment, we exposed the whole cell population to blue light. Next, we wanted to see if we could confine apoptosis to specific regions of the cell culture dish. We created a photomask, whereby light passed through only in five squares, leaving the rest of the dish non-illuminated (Fig. 5d). Since our constructs express mScarlet as transfection control (Fig. 1d), we reasoned that, by imaging the cells under the fluorescence microscope, we could detect regions with apoptosis directly in this channel without further staining. We found that both constructs led to substantial apoptosis in the illuminated areas (Fig. 5e). Light itself did not cause apoptosis (Fig. 5e, no Cre reporter control). Notably, many more cells grew in the centre of the dish than in other areas (Fig. 5e, no Cre reporter control), visually dampening the effect of the apoptosis-inducing constructs in that region.

## Discussion

In this work, we exploited an ML algorithm we recently developed, Int&in^16^, to engineer novel conditional versions of the extremely fast and efficient gp41-1 and NrdJ-1 split inteins. At the time we initiated this study, no photo-controllable gp41-1 had been reported. While our work was ongoing, Karasev and colleagues published a red-light inducible version of gp41-1 following a similar strategy as ours; that is, splitting the intein at unnatural sites^15^. They, however, did not use an algorithm to direct the selection of the split sites. Actually, this approach based on splitting an intein at unnatural sites to obtain two intein fragments with low affinity for each other that require a heterodimerization system to splice dates back to the early 2000s, when the yeast vacuolar intein VMA was artificially split to create conditional versions regulated by small molecules or light^11, 14, 46^. Currently, there is no light-inducible version of NrdJ-1. Recently, one where the external trigger is rapamycin/rapalog has been reported^9^. Moreover, NrdJ-1 has been artificially split in a previous study, but the resulting variants were either inactive or had very low activity^22^. Here, we successfully engineered novel functional NrdJ-1 split at unnatural sites.

To identify split sites in NrdJ-1 with Int&in, we used an AlphaFold-predicted structure of the protein. The crystal structure of NrdJ-1 has since been determined^47^. When we fed this structure into Int&in, several sites shifted into different categories (Supplementary Fig. 11). Of the sites we had originally selected, most retained their category, with one shifting to the inactive group. Taking the predictions based on the crystal structure into account, the precision of Int&in for NrdJ-1 increases to 62.5%.

To contextualize this value, during Int&in development we created a dataset of randomly selected split sites in gp41-1, *Npu* DnaE, and CL^16^. While 61% and 57% of these sites were experimentally active for gp41-1 and *Npu* DnaE, respectively, only 32% were active for CL. Thus, a precision of 62.5% is clearly above what would be expected from random selection. Nonetheless, Int&in performs somewhat less accurately with NrdJ-1 than with gp41-1. This can be attributed to the fact that Int&in was trained on a relatively small experimental dataset generated in *Escherichia coli* using optimal local exteins for each intein and model full-length proteins rather than split constructs. Moreover, the training readout reflected only protein reconstitution, not functional activity. Considering these limitations, the algorithm’s performance with NrdJ-1—an intein absent from the training set—is actually remarkable. Because Int&in’s predictions are agnostic to expression levels, we verified that the constructs producing no detectable Cre activity were not poorly expressed. For all split sites, both N- and C-terminal precursor proteins were visible by Western blot, ruling out low expression as the cause of absent eGFP reporter signal.

For some sites, however—such as I20:X21 in gp41-1 and F25:X26 in NrdJ-1—Int&in predicted high activity probabilities, whereas they proved inactive experimentally. Both satisfy the structural and sequence criteria expected for good sites, including surface exposure, low conservation, and strong predicted affinity of the resulting intein fragments. This highlights that the extein context can have a decisive influence on splicing efficiency.

We used a previously tested splice site for Cre^24^, which is different from those used in other optogenetic Cre designs. Interestingly, many previous studies showed functional Cre reconstitution—whereby activity is regained upon physical proximity of the Cre fragments without peptide bond formation—employing only heterodimerization systems such as the Magnets we also used^48-50^. For us, iLID or Magnets alone did not support any Cre activity (Fig. 2b and Supplementary Fig. 6). The likely explanation lies in the affinity of the Cre fragments when Cre is split at other sites than the one we selected.

It is hard to fairly and thoroughly compare the existing blue light-inducible Cre recombinases with the ones developed here, due to different cell lines, reporter assays, and illumination protocols. We can state that our light-inducible Cre recombinases feature low leakiness and good light-dark fold-changes. We envisage that they will be useful in applications in living animals, such as mice, when Cre activity needs to be triggered after several weeks of darkness during which the protein would be expressed and thus tightness is essential. In *in vitro* applications, light-inducible Cre can be used to trigger apoptosis in specific cell sub-populations, as shown as a proof of principle in a simple setup in this study. Much more sophisticated setups can be employed to pattern light in a refined manner as done by Kumar and colleagues^34^. Notably, their molecular approach to trigger apoptosis is akin to ours – regulate the expression of a constitutively active form of the human caspase-3 via a blue light-inducible synthetic transcription factor. Since the Cre recombinase irreversibly modifies the genetic information, contrary to a transcription factor that can be turned on and off with light, in our system the expression of tBID and caspase-8 is irreversibly turned on.

While the conditional gp41-1 and NrdJ-1 engineered in this study are portable, that is, they are independent of the exteins, we anticipate that optimization will be required for each case. When fusing the inteins to two protein fragments or domain with non-negligibile affinity for each other, for instance, splicing in the dark is likely to occur, something we did not observe thanks to our selection of splice site in Cre.

In conclusion, we showed that the Int&in software is a reliable tool to create conditional inteins and encourage researchers to adopt it for different inteins.

## Methods

### Prediction of intein split sites using the Int&in web server

The crystal structures of gp41-1 (PDB ID: 6QAZ) and NrdJ-1 (PDB ID: 8UBS) were uploaded on the Int&in server (freely available for academia at https://intein.biologie.uni-freiburg.de). Additionally, the structure of NrdJ-1 was predicted with AlphaFold2^21^ using its amino acid sequence as input.

### Constructions of plasmids

A DNA fragment comprising the EF1α promoter, mScarlet coding sequence, and BGH terminator (synthesized by IDT) was cloned via Gibson Assembly into the pmCherry-N1 backbone (Clontech). This is the plasmid backbone used for all subsequent cloning, expect for the Cre reporters (see below). Full-length Cre recombinase, tBID, caspase-8, gp41-1, NrdJ-1, iLID and SspB_nano were synthesized by IDT. Split variants of genes were cloned using the synthesized genes as template for the PCR. The light-inducible dimerization system pMagnet and nMagnet were amplified from Addgene plasmid #122960 (Kawano et al., Nat. Commun.). The mutations to obtain sLID from iLID, and SspB milli and micro from SspB nano were introduced by PCR followed by ligation of the linearized vectir by T4 DNA ligase cloned. (2015; Guntas et al., Nat. Methods, 2015; Zimmerman et al., Biochemistry, 2016). All plasmids were generated via Gibson Assembly using the NEBuilder HiFi DNA Assembly Kit (New England Biolabs). All plasmids cloned in this study with their features are listed in Supplementary Table S1. Except for the construct expressing mScarlet alone under the EF1α promoter, a T2A sequence (synthesized by IDT) was placed directly downstream of mScarlet and upstream of the corresponding sequence to enable co-expression by EF1α promoter. The reporter plasmid pCALVL-EGFP (Addgene #13770) was used as a template, from which tBID and caspase-8 coding sequences were inserted by replacing EGFP via AgeI/NotI (New England Biolabs) restriction cloning, yielding pCALVL-tBID and pCALVL-caspase-8 (Supplementary Table S2). PCR reactions were performed with Phusion Flash High-Fidelity PCR Master Mix (Thermo Fisher Scientific). Point mutations were introduced by primer design and incorporated by PCR-mediated plasmid linearization followed by ligation via T4 DNA ligase (New England Biolabs). Oligonucleotides were synthesized by IDT or Sigma-Aldrich. DNA sequencing was performed by Sanger sequencing. Sequencing services were provided by Microsynth. All plasmid digital maps and full sequences are available upon request.

### Cell lines and cell culture

HEK 293T (ATCC CRL-11268) and U2OS cells (kind gift of Thomas Hofmann, DKFZ, Heidelberg) were maintained in Dulbecco’s Modified Eagle Medium (DMEM; Thermo Fisher Scientific, Waltham) supplemented with 10% fetal bovine serum (FBS; Thermo Fisher Scientific), L-glutamine (Thermo Fisher Scientific), 100 U/ml penicillin and 10 mg/ml streptomycin (Thermo Fisher Scientific) at 37 °C in a humidified incubator with 5% CO_2_. Cells were passaged when reaching 80–90% confluence using 0.05% trypsin–EDTA solution (Thermo Fisher Scientific).

### Transfection

HEK 293T cells were transfected using the calcium phosphate precipitation method for experiments involving Cre-mediated expression of EGFP. Cells were seeded one day before transfection at 1.0 × 10^5 cells per well in 24-well plates, 4.0 × 10^5 cells per well in 6-well plates, or 1.8 × 10^4 cells per well in 96-well plates. Constructs were diluted in sterile water and combined with CaCl_2_ (final concentration 200 mM). The DNA–CaCl_2_ solution was added dropwise to 2× HBS buffer (50 mM HEPES, 280 mM NaCl, 1.5 mM Na_2_HPO_4_, pH 7.05) and incubated for 3 min to allow precipitate formation, followed by dropwise addition to the cells. For experiments involving Cre-mediated tBID and caspase-8 expression, HEK 293T and U2OS cells were transfected using TransitX2 (Mirus Bio) according to the manufacturer’s instructions. U2OS cells were seeded at 4.0 × 10^4 cells per well in 24-well plates one day before transfection. HEK 293T cells were seeded as described above.

### Illumination and flow cytometry analysis

Following transfection, cells were maintained in the dark for 24 h. Thereafter, samples were either kept continuously in the dark (controls) or exposed to blue light (455 nm) using a custom-built LED illumination box. For intein-based Cre-mediated recombination of the pCALVL-eGFP reporter, cells were illuminated with 13 mW/cm^2^ in pulsed cycles of 20 s light followed by 60 s dark, repeated for 6 h. After illumination, cells were incubated in the dark for an additional 18 h before harvesting by trypsinization. Recombination efficiency was quantified by flow cytometry as the ratio of GFP/mScarlet double-positive cells to mScarlet positive cells. GFP mean intensity was measured in the GFP/mScarlet double-positive population. For intein-based Cre-mediated recombination of the pCALVL-tBID and pCALVL-Caspase-8 reporters, cells were illuminated using the same pulsed light pattern (20 s light / 60 s dark) at 2.3 mW/cm^2^ for 6 h, followed by an additional 18 h incubation in the dark. Both floating and adherent cells (collected by trypsinization) were pooled and stained with Alexa Fluor 647–conjugated Annexin V (BioLegend) and Helix NP Blue (BioLegend) according to the manufacturer’s instructions. Apoptotic cells were quantified as the fraction of cells positive for either Annexin V, Helix NP, or both relative to the total cell population.

All samples were analyzed using an Attune NxT Flow Cytometer (Thermo Fisher Scientific). The gating strategy involved selecting single, viable cells from the forward and side scatter plots to exclude debris and aggregates. Fluorescence was detected in the mCherry and GFP channels for EGFP reporter assays, and in the Alexa Fluor 647 and Pacific Blue channels for apoptosis assays. Data were processed using FlowJo v10 (BD Biosciences).

### Protein extraction and western blotting

Protein samples were prepared from HEK 293T cells transfected with the relevant plasmids. After removing the culture medium, cells were gently washed with 1 ml DPBS (Thermo Fisher Scientific), and residual medium was aspirated. Cells were lysed in M-PER Mammalian Protein Extraction Reagent (Thermo Fisher Scientific) supplemented with protease inhibitor (Thermo Fisher Scientific) for 5 min at 37 °C, followed by scraping and collection. Cell lysates were clarified by centrifugation at 14,000 × g for 10 min at 4 °C, and the supernatants were collected. Samples were mixed with Fluorescent Compatible Sample Buffer (Thermo Fisher Scientific) and boiled at 95 °C for 10 min. Proteins were separated on Mini-PROTEAN TGX Precast Protein Gels (Bio-Rad) in Tris/Glycine/SDS Running Buffer (Bio-Rad), at 120 V using a PowerPac Basic power supply (Bio-Rad). Proteins were transferred to PVDF membranes using the Trans-Blot Turbo Transfer System (Bio-Rad). Membranes were blocked at room temperature for 1 h with Blocker™ FL Fluorescent Blocking Buffer (Thermo Fisher Scientific), and incubated overnight at 4 °C with primary antibodies rat anti-FLAG (BioLegend, 1:1000) and rabbit anti-β-tubulin (Cell signaling, 1:10000). After three washes with TBS-T buffer (20 mM Tris-HCl, 150 mM NaCl, 0.05% Tween-20, pH 7.4), membranes were incubated for 1 h at room temperature in the dark with secondary antibodies Alexa Fluor 488-conjugated goat anti-rat IgG (Invitrogen, 1:1000) and Alexa Fluor 647-conjugated donkey anti-rabbit IgG (Invitrogen, 1:1000). Membranes were washed three times with TBS-T and imaged using a Typhoon scanner (Amersham, Cytiva). For detection via bioluminescence, membranes were incubated with Restore Western Blot Stripping Buffer (Thermo Fisher Scientific) at 37 °C for 15 min, washed three times with TBS-T, and processed according to the Nano-Glo® HiBiT Blotting System protocol (Promega). HiBiT signals were detected using a Fusion imaging system (Vilber). Image analysis was performed using Fiji (ImageJ; National Institutes of Health, Bethesda, MD, USA).

### Light titration assay using the Diya illumination device

HEK 293T cells were seeded in a black 96-well plate with transparent bottom (μ-Plate 96 Well Black, ibidi) one day before transfection. During the illumination, plates were placed in a custom-made black 96-well holder that aligned each well with an individual optical channel, ensuring that light from the underlying LED reached only the corresponding well. Blue light (455 nm) illumination was performed using the Diya illumination device^32^. The light intensity was set to start at 1.8 mW/cm^2^ (setting = 255) and then gradually decreased across the plate in a well-to-well gradient down to 0 mW/cm^2^ (setting = 0), with increments of 30 in the Diya setting. The illumination program consisted of pulsed of blue light (20 s) followed by a dark phase (40 s), repeated for 12 h. After illumination, cells were harvested and analyzed by flow cytometry as described above.

### Patterned illumination with a photomask

Patterns were printed on paper and used as template to cut the same shape onto an aluminum foil, which was then attached onto a glass support. This custom-made photomask (allowing light to go through only where the aluminum foil was cut away) was placed onto the cell culture dish. 24 h post-transfection, culture dishes with the photomask were placed in the custom-built LED illumination box. For the experiment with the eGFP reporter, cells were plated in 6-well plates and illuminated with 455 nm light at an intensity of 2.3 mW/cm^2^ for 2 h. For experiments with the pCALVL-tBID/Caspase-8 reporter and control, cells were plated in 24-well plates and illuminated with the same light regime for 6 h. Cells were kept in the dark for 24 h. Afterwards, images were acquired using a Lionheart FX Automated Microscope (Agilent BioTek) in brightfield, GFP, and Texas Red channels.

### Statistics and reproducibility

Unless otherwise stated, all experiments were conducted independently three times, and consistent results were obtained across replicates. The sample size was not predetermined using statistical methods but was chosen based on prior experience and established experimental practices. No data were excluded from analysis. Experiments were not randomized. The investigator was not blinded to group allocation during experimentation or outcome assessment; however, samples were allocated randomly. Details on sample sizes (biological replicates, *n*), statistical analyses, and significance levels are provided in the figure legends. Statistical significance was assessed using unpaired, two-tailed Student’s *t*-tests with multiple-comparison correction according to the Holm–Šidák procedure. Statistical analyses were performed using GraphPad Prism v 10.12 (GraphPad Software Inc.).

## Acknowledgments

We thank Timothy Curtis Shoyer for critical reading of the manuscript, Mehmet Ali Öztürk and Franziska Schneider-Warme for discussions and feedback during the course of this study, and Mustafa Khammash for the generous donation of the Diya illumination device. We acknowledge the scientific and technical support of the Signalling Factory & Robotics of the Albert-Ludwigs-University Freiburg, especially Pavel Salavei, for help with the flow cytometer, and the excellent support in data recording and analysis. We also thank the Weber group for providing the apparatus for the experiment with the photomask. This study was part of SFB1425, funded by the Deutsche Fourschungsgemeinschaft (DFG, German Research Foundation) – Project #422681845. ChatGPT (OpenAI) was used to improve grammar and flow.

## Author contributions

J.L. performed all experiments, analysed data and prepared figures. B.D.V. conceived the project, supervised the study, analysed data, provided funding and wrote the manuscript. J.L. read and approved the manuscript.

## Data availability

The Cre constructs used in Figs. 4 and 5 are available from Addgene. Other constructs are available upon request.

