## Supplementary Figures 1-12 for "Machine learning–guided engineering of conditional split inteins for regulated protein splicing in mammalian cells"

for

Supplementary Figures 1-12

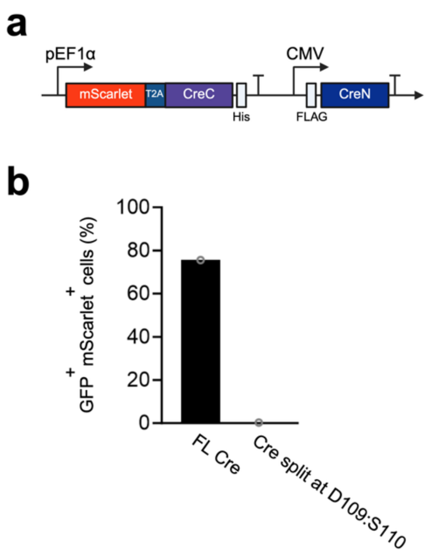


**Supplementary Fig. 1 | Cre split at D109:S110 does not functionally reconstitute on its own.**

**a** Schematic representation of the split Cre construct. T2A, ribosome skipping sequence of the Thosea asigna virus. **b** Bar graph showing the percentage of HEK 293T cells transfected with the indicated constructs (mScarlet-positive) in which Cre successfully recombined the reporter DNA (GFP-positive) measured via flow cytometry. This experiment was done once. FL Cre, full-length Cre expressed from the strong, constitutive CMV promoter. Created with BioRender.

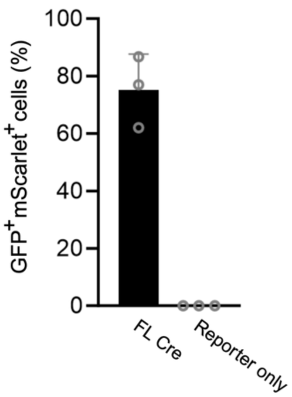


**Supplementary Fig. 2 | The STOP cassette in the reporter construct is stable in the absence of Cre recombinase.**

Bar graph showing the percentage of HEK 293T cells transfected with the indicated constructs (mScarlet-positive) showing also GFP fluorescence, indicating recombination of the reporter DNA, measured via flow cytometry. Values represent mean ± SD of n = 3 biologically independent experiments. Individual data points are shown as open circles. FL Cre, full-length Cre expressed from the strong, constitutive CMV promoter. Created with BioRender.

**a**

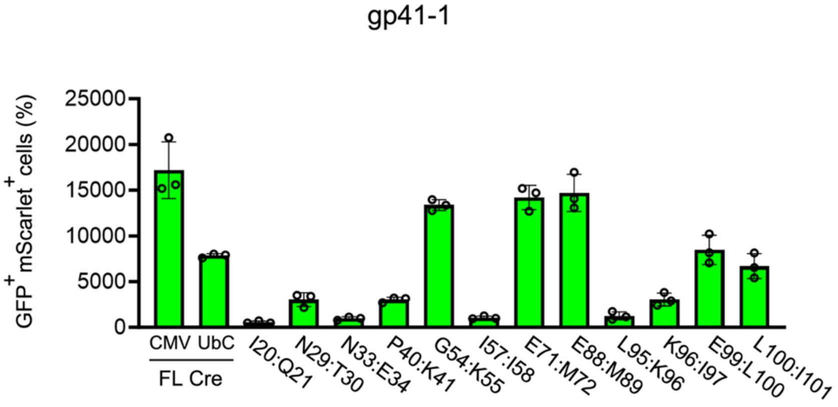


**b**

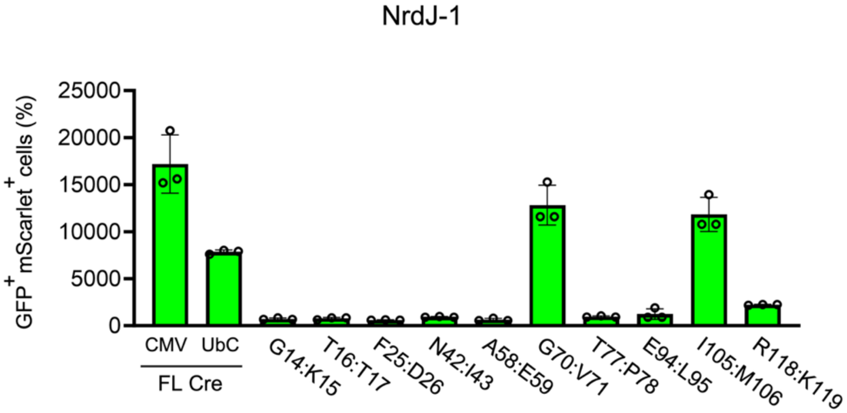


**Supplementary Fig. 3 | Quantification of mean GFP intensity in HEK 293T cells transfected with split Cre fused to gp41-1 and NrdJ-1 split at unnatural sites.**

**a,b** Bar graphs showing the mean GFP fluorescence of HEK 293T cells transfected with the indicated gp41-1 (**a**) or NrdJ-1 (**b**) constructs (mScarlet-positive) in which Cre successfully recombined the reporter DNA (GFP-positive) measured via flow cytometry. Values represent mean ± SD of n = 3 biologically independent experiments. Individual data points are shown as open circles. FL Cre, full-length Cre expressed from either the strong constitutive CMV promoter or the weak UbC promoter. Created with BioRender.

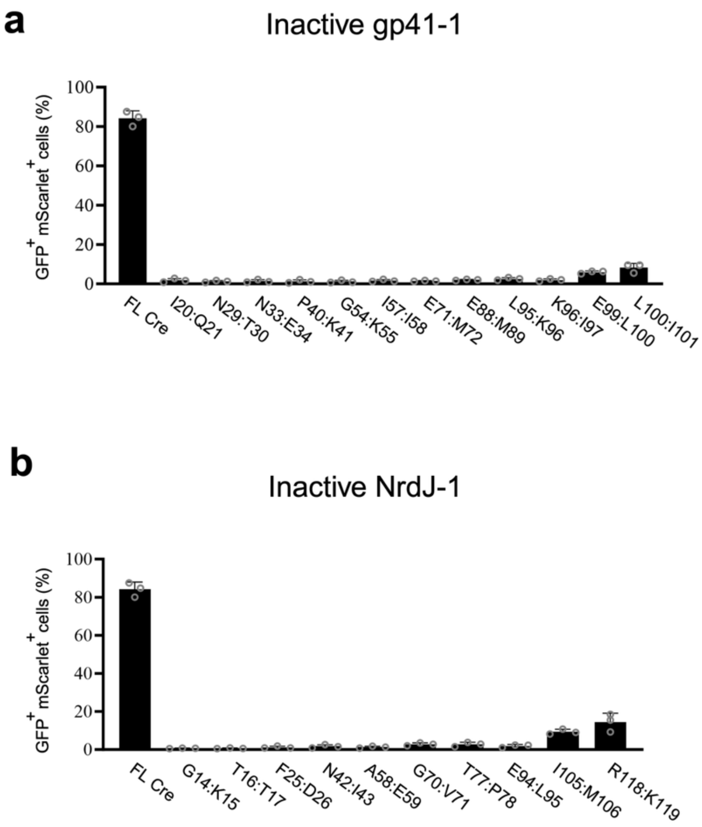


**Supplementary Fig. 4 | Cre activity is dependent on intein-mediated protein splicing.**

**a,b** Bar graphs showing the percentage of HEK 293T cells transfected with the indicated constructs (mScarlet-positive) in which Cre successfully recombined the reporter DNA (GFP-positive) measured via flow cytometry. gp41-1 and NrdJ-1 were inactivated by mutating the conserved N-terminal cysteine and C-terminal asparagine to alanine. Values represent mean ± SD of n = 3 biologically independent experiments. Individual data points are shown as open circles. FL Cre, full-length Cre expressed from the strong, constitutive CMV promoter. Created with BioRender.

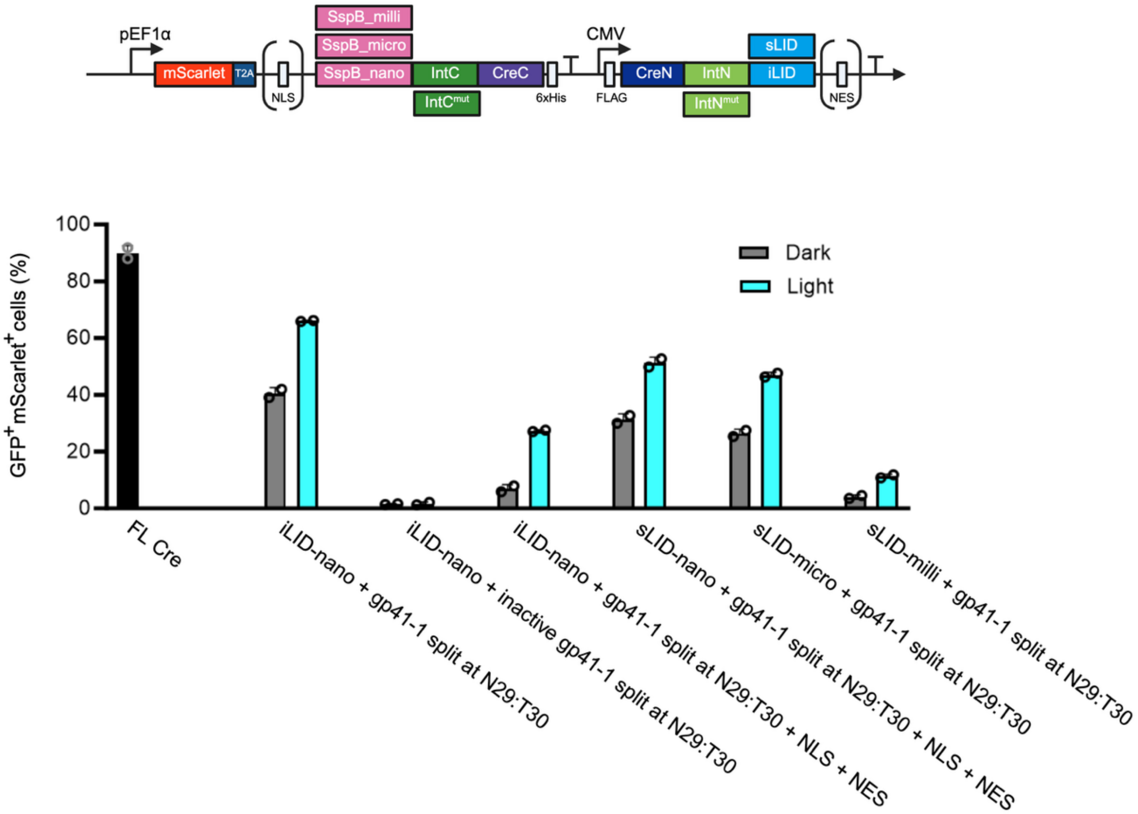


**Supplementary Fig. 5 | Optimization of iLID-based constructs and assessment of intein-mediated splicing contribution to Cre recombinase activity.**

Upper panel, schematic representation of the constructs. The elements in parentheses (nuclear import signal (NLS); nuclear export sequence (NES)) are present in some of the constructs only. One of the constructs contains inactive gp41-1 split at N29:T30. The intein was inactivated by mutating the conserved N-terminal cysteine and C-terminal asparagine to alanine. T2A, ribosome skipping sequence of the Thosea asigna virus. Lower panel, bar graph showing the percentage of HEK 293T cells transfected with the indicated constructs (mScarlet-positive) in which Cre successfully recombined the reporter DNA (GFP-positive) measured via flow cytometry. Values represent mean ± SD of n = 2 biologically independent experiments. Individual data points are shown as open circles. FL Cre, full-length Cre expressed from the strong, constitutive CMV promoter. Cells were either kept in the dark the whole time or illuminated for 6 hours with blue light (13 mW/cm²) in short pulses (20 s light / 60 s dark) followed by additional incubation in the dark for 18 hours. Cells expressing FL Cre were kept in the dark. Created with BioRender.

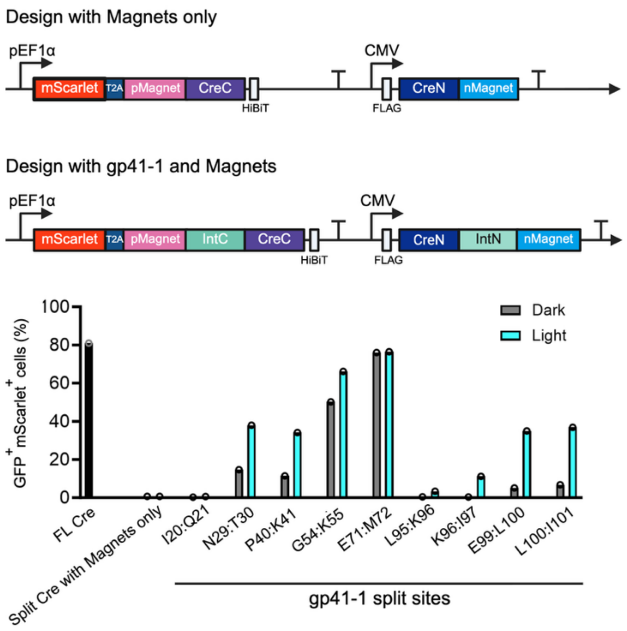


**Supplementary Fig. 6 | Magnets-based design of conditional gp41-1 improves Cre performance.**

Upper and middle panels, schematic representation of the constructs. T2A, ribosome skipping sequence of the Thosea asigna virus. Lower panel, bar graph showing the percentage of HEK 293T cells transfected with the indicated constructs (mScarlet-positive) in which Cre successfully recombined the reporter DNA (GFP-positive) measured via flow cytometry. Values represent single measurements (n = 1). FL Cre, full-length Cre expressed from the strong, constitutive CMV promoter. Cells were either kept in the dark the whole time or illuminated for 6 hours with blue light (13 mW/cm²) in short pulses (20 s light / 60 s dark) followed by additional incubation in the dark for 18 hours. Cells expressing FL Cre were kept in the dark. Created with BioRender.

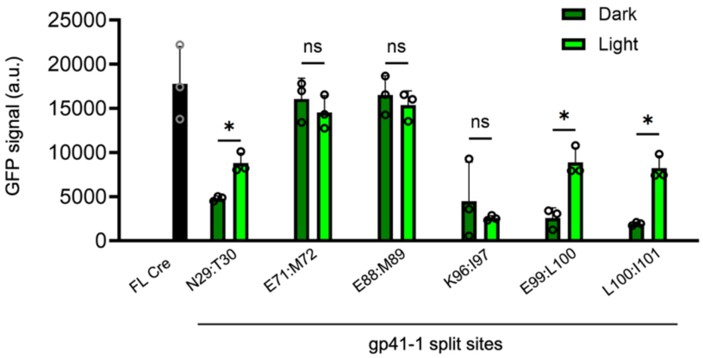


**Supplementary Fig. 7 | The mean GFP fluorescence is different in illuminated and non-illuminated samples for some of the gp41-1 constructs.**

Bar graph showing the mean GFP fluorescence of HEK 293T cells transfected with the indicated constructs (mScarlet-positive) in which Cre successfully recombined the reporter DNA (GFP-positive) measured via flow cytometry. Values represent mean ± SD of n = 3 biologically independent experiments. Individual data points are shown as open circles. Statistical significance was determined by unpaired, two-tailed Student's *t*-tests, with multiple-comparison correction according to the Holm-Šidák procedure. *, p-value $\leq 0.05$; ns, p-value $> 0.05$. FL Cre, full-length Cre expressed from the strong, constitutive CMV promoter. Cells were either kept in the dark the whole time or illuminated for 6 hours with blue light (13 mW/cm²) in short pulses (20 s light / 60 s dark) followed by additional incubation in the dark for 18 hours. Cells expressing FL Cre were kept in the dark. Created with BioRender.

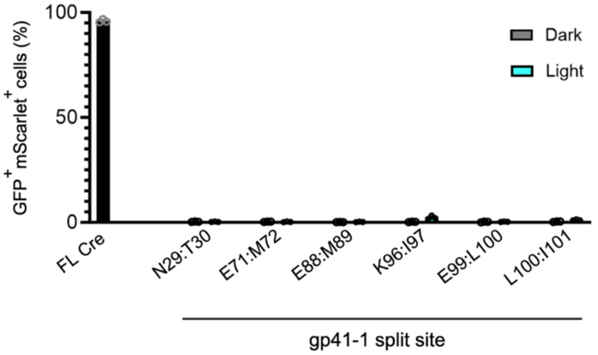


**Supplementary Fig. 8 | Cre activity is dependent on gp41-1-mediated protein *trans*-splicing.**

Bar graph showing the percentage of HEK 293T cells transfected with indicated constructs (mScarlet-positive) in which Cre successfully recombined the reporter DNA (GFP-positive) measured via flow cytometry. gp41-1 in all split constructs was inactivated by mutating the conserved N-terminal cysteine and C-terminal asparagine to alanine. Values represent mean ± SD of n = 3 biologically independent experiments. Individual data points are shown as open circles. FL Cre, full-length Cre expressed from the strong, constitutive CMV promoter. Cells were either kept in the dark the whole time or illuminated for 6 hours with blue light (13 mW/cm²) in short pulses (20 s light / 60 s dark) followed by additional incubation in the dark for 18 hours. Cells expressing FL Cre were kept in the dark. Created with BioRender.

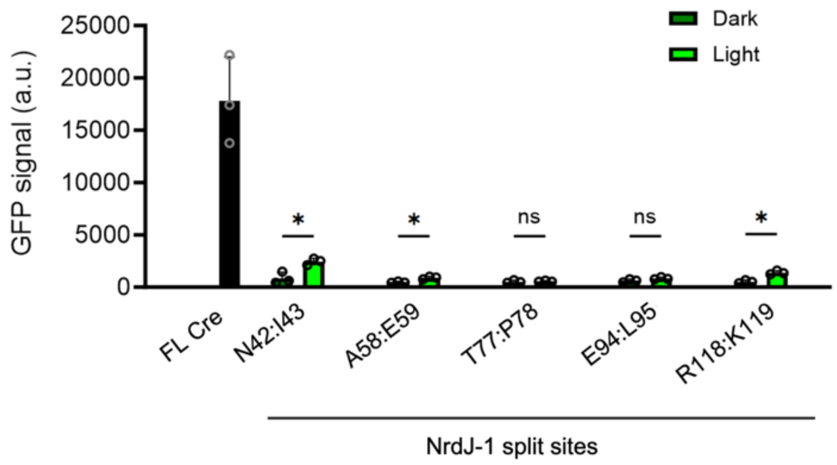


**Supplementary Fig. 9 | HEK 293T cells transfected with the engineered, conditional NrdJ-1 variants split at N42:I43 and R118:K119 have higher mean GFP fluorescence when illuminated.**

Bar graph showing the mean GFP fluorescence of cells transfected with the indicated constructs (mScarlet-positive) in which Cre successfully recombined the reporter DNA (GFP-positive) measured via flow cytometry. Values represent mean ± SD of n = 3 biologically independent experiments. Individual data points are shown as open circles. Statistical significance was determined by unpaired, two-tailed Student's *t*-tests, with multiple-comparison correction according to the Holm-Šidák procedure. *, p-value $\leq 0.05$; ns, p-value $> 0.05$. FL Cre, full-length Cre expressed from the strong, constitutive CMV promoter. Cells were either kept in the dark the whole time or illuminated for 6 hours with blue light (13 mW/cm²) in short pulses (20 s light / 60 s dark) followed by additional incubation in the dark for 18 hours. Cells expressing FL Cre were kept in the dark. Created with BioRender.

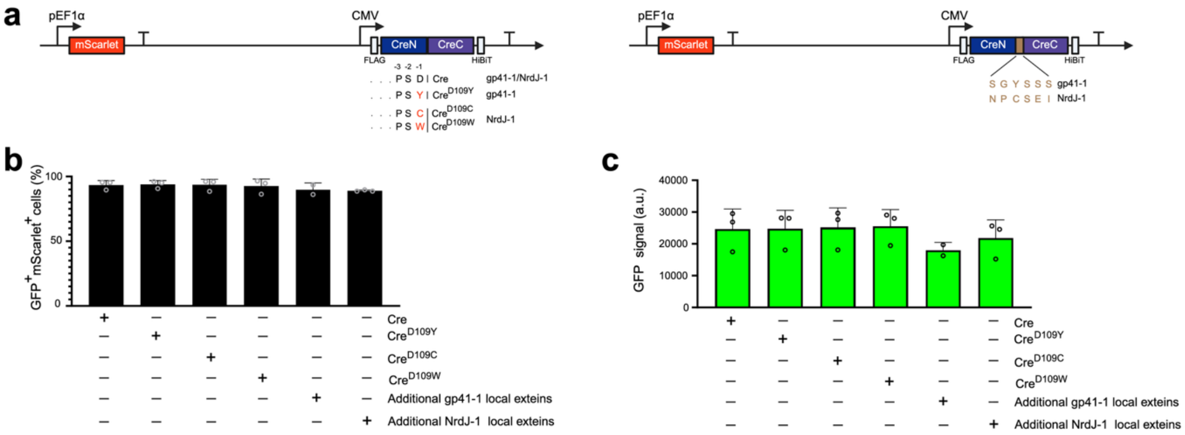


**Supplementary Fig. 10 | Mutation of D109 or insertion of amino acids in Cre recombinase does not affect its activity.**

**a** Schematic representation of the constructs. **b,c** Bar graphs showing the percentage (**b**) or mean GFP fluorescence (**c**) of cells transfected with the indicated constructs (mScarlet-positive) in which Cre successfully recombined the reporter DNA (GFP-positive) measured via flow cytometry. Values represent mean ± SD of n = 3 biologically independent experiments. Individual data points are shown as open circles. In this experiment, Cre is not split; nonetheless, for easier comparison with split-Cre constructs and to visualize the location of the local extein-based linker, the protein is depicted as N- and C-terminal fragments. Created with BioRender.

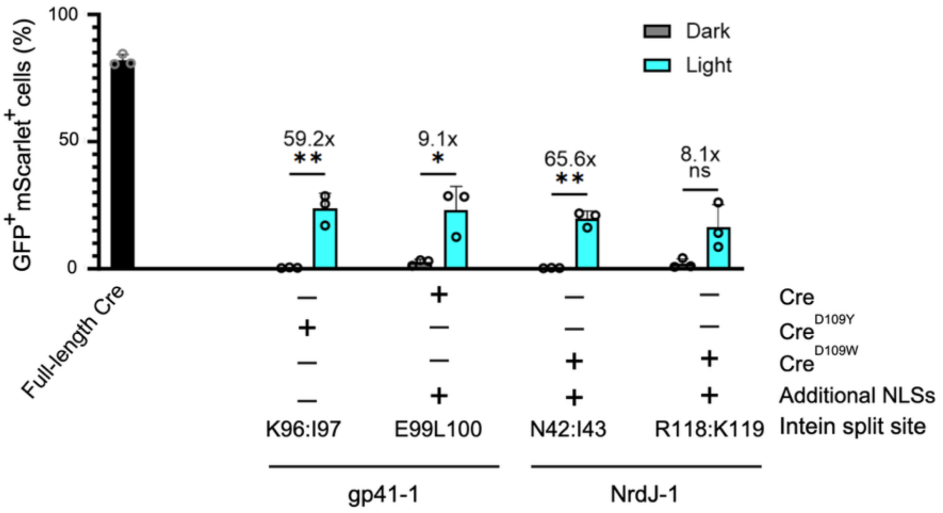


**Supplementary Fig. 11 | The intein-controlled light-inducible Cre is functional in U2OS cells.**

Bar graph showing the percentage of cells transfected with the indicated constructs (mScarlet-positive) in which Cre successfully recombined the reporter DNA (GFP-positive) measured via flow cytometry. Values represent mean ± SD of n = 3 biologically independent experiments. Individual data points are shown as open circles. Full-length Cre was expressed from the strong, constitutive CMV promoter. Cells were either kept in the dark the whole time or illuminated for 6 hours with blue light (2.3 mW/cm²) in short pulses (20 s light / 60 s dark) followed by additional incubation in the dark for 18 hours. Cells expressing FL Cre were kept in the dark. Cells expressing FL Cre were kept in the dark. Statistical significance was determined by unpaired, two-tailed Student's *t*-tests, with multiple-comparison correction according to the Holm-Šidák procedure. **, p-value ≤ 0.01; *, p-value ≤ 0.05; ns, p-value > 0.05. The light-dark fold change is indicated above each construct. Created with BioRender.

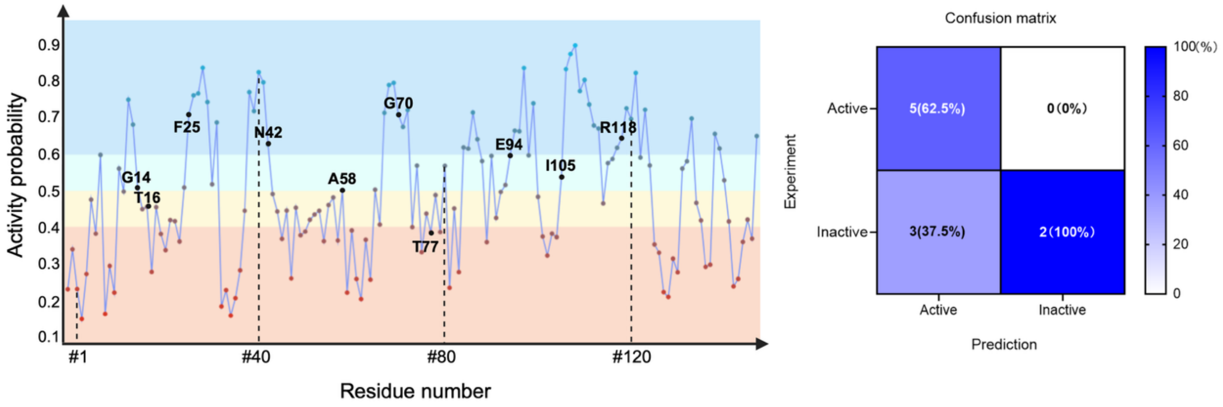


**Supplementary Fig. 12 | Feeding the NrdJ-1 crystal structure to Int&in alters the algorithm output.**

Left panel, output of the Int&in algorithm for NrdJ-1 when using the crystal structure (PDB: 8UBS). The four categories are: active split sites ($p \geq 0.5$), inactive split sites ($p < 0.5$), active with high probability ($p \geq 0.6$), inactive with high probability ($p < 0.4$). The sites selected for experimental testing are indicated in black. Numbering starts at the third amino acid, as the first two residues correspond to extein sequences present in the crystal structure. Int&in is accessible at https://intein.biologie.uni-freiburg.de. Right panel, confusion matrix illustrating the performance of the Int&in prediction algorithm. Columns correspond to predicted classes, and rows to experimentally-determined classes. Percentages are normalized by column, indicating the proportion of correct predictions within each class. The intein was considered active if the percentage of double-positive cells was above 4%.
